# The Pathfinder plasmid toolkit for genetically engineering newly isolated bacteria enables the study of *Drosophila*-colonizing *Orbaceae*

**DOI:** 10.1101/2023.02.15.528778

**Authors:** Katherine M. Elston, Laila E. Phillips, Sean P. Leonard, Eleanor Young, Jo-anne C. Holley, Tasneem Ahsanullah, Braydin McReynolds, Nancy A. Moran, Jeffrey E. Barrick

## Abstract

Toolkits of plasmids and genetic parts streamline the process of assembling DNA constructs and engineering microbes. Many of these kits were designed with specific industrial or laboratory microbes in mind. For researchers interested in non-model microbial systems, it is often unclear which tools and techniques will function in newly isolated strains. To address this challenge, we designed the Pathfinder toolkit for quickly determining the compatibility of a bacterium with different plasmid components. Pathfinder plasmids combine three different broad-host-range origins of replication with multiple antibiotic resistance cassettes and reporters, so that sets of parts can be rapidly screened through multiplex conjugation. We first tested these plasmids in *Escherichia coli*, a strain of *Sodalis praecaptivus* that colonizes insects, and a *Rosenbergiella* isolate from leafhoppers. Then, we used the Pathfinder plasmids to engineer previously unstudied bacteria from the family *Orbaceae* that were isolated from several fly species. Engineered *Orbaceae* strains were able to colonize *Drosophila melanogaster* and could be visualized in fly guts. *Orbaceae* are common and abundant in the guts of wild-caught flies but have not been included in laboratory studies of how the *Drosophila* microbiome affects fly health. Thus, this work provides foundational genetic tools for studying new host-associated microbes, including bacteria that are a key constituent of the gut microbiome of a model insect species.

**IMPORTANCE:** To fully understand how microbes have evolved to interact with their environments, one must be able to modify their genomes. However, it can be difficult and laborious to discover which genetic tools and approaches work for a new isolate. Bacteria from the recently described *Orbaceae* family are common in the microbiomes of insects. We developed the Pathfinder plasmid toolkit for testing the compatibility of different genetic parts with newly cultured bacteria. We demonstrate its utility by engineering *Orbaceae* strains isolated from flies to express fluorescent proteins and characterizing how they colonize the *Drosophila melanogaster* gut. *Orbaceae* are widespread in *Drosophila* in the wild but have not been included in laboratory studies examining how the gut microbiome affects fly nutrition, health, and longevity. Our work establishes a path for genetic studies aimed at understanding and altering interactions between these and other newly isolated bacteria and their hosts.

## INTRODUCTION

Researchers have isolated and sequenced many new microbes from different ecosystems and from diverse plant and animal hosts. To characterize these microbes and study how they interact with their physical environments and with other organisms, one needs genetic tools. However, most described bacterial species have never been genetically manipulated (1–3). The primary obstacle in many cases is likely that the requisite resources and know-how for microbial genetic engineering are not easily accessible to researchers who encounter non-model microbes.

Toolkits of genetic parts have been developed for molecular microbiology and synthetic biology. These kits, such as the modular cloning (MoClo) and Standard European Vector Architecture (SEVA) toolkits (4, 5), aim to be flexible and comprehensive. Their collections of interchangeable parts include promoters with a range of different expression levels and multiple reporter genes and plasmid backbones that can be combined to assemble a genetic construct of interest (4, 6–11). While these systems facilitate complex genetic engineering tasks in well-studied laboratory and industrial bacteria, such as *Escherichia coli* and *Pseudomonas putida*, there are still gaps with respect to their applicability to all bacteria. For example, they may only have a few antibiotic resistance cassettes or rely on plasmids that replicate only in specific species. Researchers may also find assembling new plasmids according to the schemes in these kits daunting and overly cumbersome if all they want to achieve are basic tasks like expressing a single protein in a new bacterial species.

Fluorescent protein expression alone is often enough to investigate aspects of host-microbe interactions, such as bacterial localization, or to track a strain of interest within a microbiome or environmental community. However, even this rudimentary genetic modification can be challenging (1, 3, 12). A reasonable first step is to start with a broad-host-range plasmid that has been reported to replicate in diverse bacteria, but one must still empirically test whether one of these plasmids is compatible with each new species (13). Additionally, electroporation or chemical treatments to transform plasmids into cells do not work in all bacteria (14, 15). Developing these techniques through trial and error can be frustrating and time-consuming, particularly when one does not know if the plasmid being used will successfully replicate after transformation. Conjugation is often a more reliable method for delivering DNA to non-model bacteria (2, 16), and it has been incorporated into several kits that focus on engineering a wider range of bacteria, such as the Bacterial Expression Vector Archive (BEVA) (9), the bee microbiome toolkit (BTK) (6), and the Proteobacteria toolbox (8). The wide phylogenetic distribution of natural conjugative and transmissible broad-host-range plasmids suggests that this approach should work for many bacterial species (17–19).

Although *Drosophila melanogaster* has been a model organism for genetics for over a century, research focused on its gut microbiome is a relatively new field (20–22). Laboratory studies have focused primarily on *Acetobacter* and *Lactobacillus* species (23–25), which make up only a fraction of the microbiome that is normally present in wild *Drosophila* (26). A large percentage of the natural *Drosophila* microbiome is composed of bacteria in the recently-described *Orbaceae* family (27, 28). *Orbaceae* are prevalent in a wide variety of insects (28–31) and are observed in 16S rRNA gene surveys of populations of laboratory-reared and wild flies of different species, including *D. melanogaster* (26, 29, 32–34). How *Orbaceae* colonize and interact with their hosts is relatively unexplored despite how prevalent they are in insect microbiomes.

Here we describe the Pathfinder plasmid system, a simple and robust toolkit for engineering newly cultured bacteria. First, we show how multiplex conjugation with defined subsets of Pathfinder plasmids can be used to quickly determine the compatibility of bacteria with different genetic parts. Then, we then use the Pathfinder plasmids to engineer recently cultured *Orbaceae* isolates from flies and characterize how they colonize the *D. melanogaster* gut.

## RESULTS

### Pathfinder plasmid toolkit design

An overview of the Pathfinder plasmid design and procedure is shown in Figure 1A and Table 1. Plasmids pSL1, pSL9, and pSL25 have 3 different origins of replication (RSF1010, pBBR1, and RP4) paired with 3 different reporters (red chromoprotein, RCP; E2-Crimson, E2C; and blue chromoprotein, BCP), respectively, along with kanamycin resistance. The reporter genes are all expressed from the broad-host-range CP25 promoter. Plasmids pSL1-7 all have an RSF1010 origin and red chromoprotein expression with one of 7 different antibiotic resistances (in order): kanamycin (Kan^R^), spectinomycin (Spec^R^), gentamicin (Gent^R^), chloramphenicol (Cam^R^), erythromycin (Ery^R^), tetracycline (Tet^R^), and ampicillin/carbenicillin (Carb^R^). All Pathfinder plasmids are also BsaI dropout vectors (Type 8 parts) compatible with stage 1 of the Golden Gate assembly scheme used by the yeast and bee microbiome toolkits (6, 35). Thus, the Pathfinder plasmids can be readily reconfigured to convey and express DNA sequences other than the included reporter genes. For example, we created pSL1-GFP to express GFP instead of RCP from the same backbone as pSL1 in this way.

**Table 1.** Pathfinder plasmids

| Plasmid | Addgene accession | Origin | Origin source | Reporter | Reporter source | Antibiotic resistance gene | Resistance source |
| --- | --- | --- | --- | --- | --- | --- | --- |
| pSL1 | 180422 | RSF1010 | pBTK402 (6) | RCP (mRFP1) | BBa_E1010 (64) | Kan (*aphA-1*) | pBTK402 (6) |
| pSL1-GFP<sup>a</sup> | 180420 | RSF1010 | pBTK402 (6) | GFP | pBTK520 (6) | Kan (*aphA-1*) | pBTK402 (6) |
| pSL2<sup>b</sup> | 190998 | RSF1010 | pBTK402 (6) | RCP (mRFP1) | Bba_E1010 (64) | Spec (*aadA*) | pBTK403 (6) |
| pSL3 | 190999 | RSF1010 | pBTK402 (6) | RCP (mRFP1) | Bba_E1010 (64) | Gent (*aacC1*) | pKNOCK-Gm (65) |
| pSL4<sup>b</sup> | 191000 | RSF1010 | pBTK402 (6) | RCP (mRFP1) | Bba_E1010 (64) | Cam (*cat1*) | pYTK001 (35) |
| pSL5<sup>b</sup> | 191001 | RSF1010 | pBTK402 (6) | RCP (mRFP1) | Bba_E1010 (64) | Ery (*ermB*) | pMSP353 5 (66) |
| pSL6 | 191002 | RSF1010 | pBTK402 (6) | RCP (mRFP1) | Bba_E1010 (64) | Tet (*tetC*) | pBMTBX-4 (67) |
| pSL7 | 191003 | RSF1010 | pBTK402 (6) | RCP (mRFP1) | Bba_E1010 (64) | Carb (*tem116*) | pBTK401 (6) |
| pSL9 | 191004 | pBBR1 | pBBR1MC S-2 (43) | E2C | pBTK570 (6) | Kan (*aphA-1*) | pBTK402 (6) |
| pSL25 | 191005 | RP4 | pTD-C_sYFPT winStrep (68) | BCP (amilCP) | amilCP chromoprotein (69) | Kan (*aphA-1*) | pBTK402 (6) |

<sup>a</sup>Plasmid is not compatible with Golden Gate assembly.
<sup>b</sup>Plasmid contains a duplication of one BsaI site and the adjacent CP25 promoter and ribosome binding site for RCP. It is still compatible with Golden Gate assembly.

**FIG 1.**
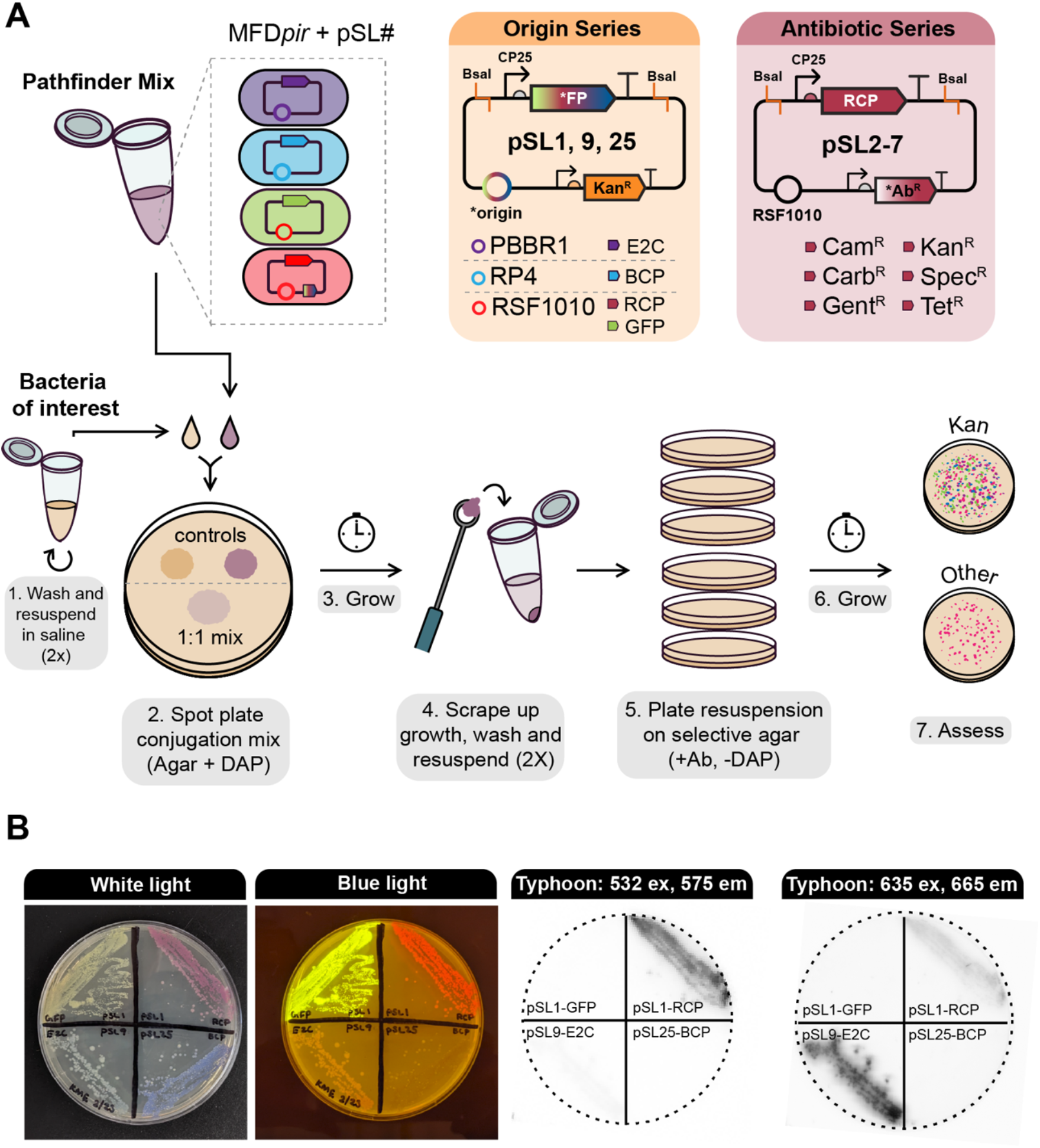
The Pathfinder plasmid system. (A) Plasmid maps and workflow for multiplex conjugation into a bacterium of interest. (B) Visualizing Pathfinder reporters. The same agar plate containing streaks of *E. coli* MFD*pir* donor strains, each with a plasmid expressing a different fluorescent protein or chromoprotein is shown in all images. The leftmost panel shows the plate under white light, the second shows the plate on a blue light transilluminator, and the last two panels show the plate imaged using a Typhoon 9500 FLA system with two different excitation and emission settings.

To confirm the functionality of the Pathfinder plasmids, we performed an initial test with *E. coli*. All plasmids except pSL5 (RSF1010, RCP, Ery^R^) were transformed into the *E. coli* donor strain MFD*pir* (36), then combined equally into a mix that could be frozen down and thawed as needed for conjugation. We were able to recover *E. coli* transconjugants using this mix for every plasmid except for pSL3 (RSF1010, RCP, Gent^R^). Colonies of *E. coli* cells containing Pathfinder plasmids expressing each of the three reporters or GFP can be identified by eye (Fig. 1B). Fluorescence can also be used to distinguish colonies with different plasmids from one another (Fig. 1B), which might be useful if the markers are expressed at lower levels in other species. To demonstrate functionality in a more distantly-related bacterium, we tested conjugation of the Pathfinder mix into *Sodalis praecaptivus* HS^T^, a human wound bacterial isolate previously shown to colonize weevils and tsetse flies (37–39). As with *E. coli*, we were able to recover transconjugants for every plasmid except for pSL3. This negative result may be due to inherent gentamicin resistance of the MFD*pir* strain interfering with conjugation. We also established that Pathfinder plasmids function in a new *Rosenbergiella* bmE01 strain we isolated from *Empoasca* leafhoppers. For bmE01, we tested only pSL1-GFP and pSL7 and successfully isolated transconjugants for both.

### Pathfinder plasmids function in recently isolated fly symbionts

We then applied the Pathfinder plasmid system to a set of bacteria that we isolated from wild flies (members of order Diptera), along with an isolate, *Orbus hercynius* CN3, collected by Volkmann *et al*. (27), which likely originates from a non-*Drosophila* dipteran species breeding in boar feces. Based on phylogenies constructed using 16S rRNA genes, all of these isolates are closely related and belong to the *Orbaceae* family within the *Gammaproteobacteria*, which includes symbionts of bees and other insects (Fig. 2A).

**FIG 2.**
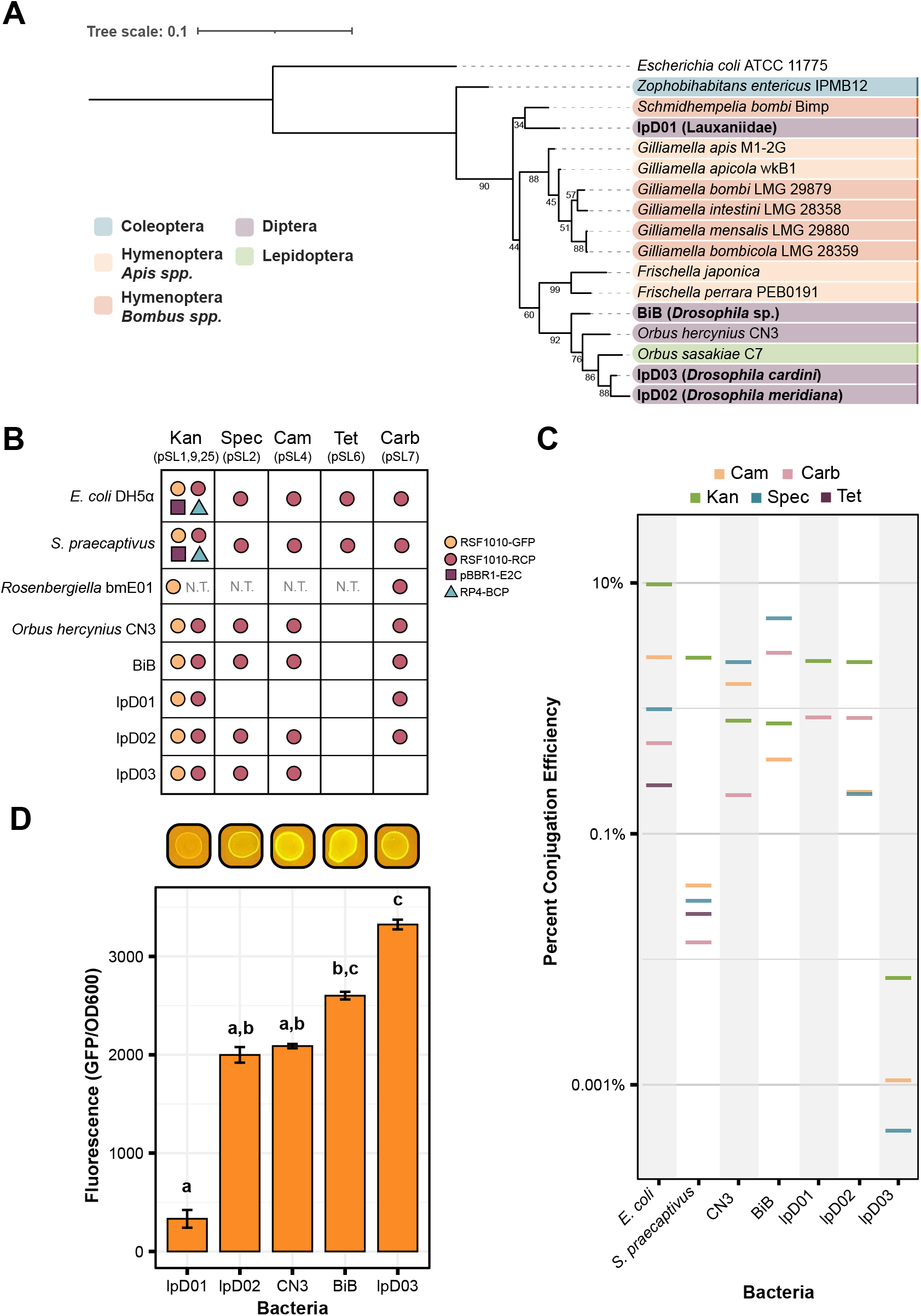
Recently isolated *Orbaceae* strains can be engineered with the Pathfinder plasmid system. (A) 16S rRNA gene sequence phylogeny showing the relationship between the isolated species in this paper (in bold) and other members of the family *Orbaceae*. Strains are color-coded based on the taxonomic order of their original insect host. For each *Orbaceae* strain first reported in this study the closest taxonomic identifier that could be established for the host is shown in parentheses. Bootstrap values are depicted next to their respective branches. (B) Table depicting the compatibility of the bacteria in this study with each of the Pathfinder plasmids. Compatibility is defined as isolation and verification of at least a single transconjugant colony for each plasmid. Dots are used to represent compatibility and are color-coded according to the origin of replication and reporter gene on the plasmid. (C) The efficiency of conjugation under each of the antibiotic conditions in the study is plotted as a percentage of transconjugants relative to growth on nonselective media. Data are plotted on a log scale. Except in the Kan condition, where multiple plasmids may be present as shown in (B), all plasmids are RSF1010-RCP. (D) Bar chart showing the level of GFP expression for each strain engineered with pSL1-GFP (RSF1010-GFP) normalized to an OD600 reading for that same strain. Images above the chart show the appearance of each strain on a blue light transilluminator. Letters above each bar designate groups that are significantly different at the *p* < 0.01 level calculated using Dunn’s test with Bonferroni correction.

We were able to successfully conjugate Pathfinder plasmids into each of the *Orbaceae* (Fig. 2B). In terms of origin of replication compatibility, we only observed conjugation with the pSL1 and pSL1-GFP plasmids that contain the RSF1010 origin. The pBBR1 and RP4 origin plasmids were absent from our *Orbaceae* transconjugant plates. For the antibiotic resistance panel, we achieved conjugation with all plasmids other than pSL6 (TetR) in most strains. Differences arose when strains had elevated levels of intrinsic resistance to an antibiotic (Table 2). For instance, lpD01 was highly resistant to Spec and Cam which prevented us from isolating pSL2 and pSL4 transconjugants, respectively, because untransformed cells grew on these selective plates. Rates of conjugation with the Pathfinder plasmids were similar across strains and comparable to the rates observed for *E. coli* for many combinations of plasmids and strains (Fig. 2C). The overall average conjugation efficiency for all the *Orbaceae* strains was 1.4 ± 0.02%. BiB had the highest average conjugation efficiency (2.3 ± 0.1%), while lpD03 had the lowest (0.0026 ± 0.0002%). Despite the lower conjugation efficiencies observed for some *Orbaceae* strains, we demonstrated that each of these newly isolated bacteria can be engineered with several plasmids from the Pathfinder series.

**Table 2.** Antibiotic concentrations used in this study *^a^*

|  |  | CARB | CAM | KAN | SPEC | TET |
| --- | --- | --- | --- | --- | --- | --- |
| *DH5α* | MIC | 100 | 20 | 50 | 60 | 10 |
|  | Pathfinder | 100 | 20 | 50 | 60 | 10 |
| *Sodalis praecaptivus* | MIC | 200 | 6.25 | 6.25 | 50 | 12.5 |
|  | Pathfinder | 200 | 6.25 | 6.25 | 50 | 12.5 |
| bmE01 | MIC | 400 | 25 | 200 | 400 | 6.25 |
|  | Pathfinder | **\*800** | **\*50** | **\*400** | **\*800** | **\*12.5** |
| *Orbus hercynius* CN3 | MIC | 800 | 6.25 | 25 | 25 | 6.25 |
|  | Pathfinder | 800 | 6.25 | 25 | **\*100** | **\*3** |
| BiB | MIC | 25 | 6.25 | 400 | 100 | 6.25 |
|  | Pathfinder | 25 | 6.25 | 400 | **\*200** | **\*3** |
| IpD01 | MIC | 6.25 | 400 | 12.5 | 400 | 200 |
|  | Pathfinder | **\*25** | 400 | **\*50** | **\*800** | 200 |
| IpD02 | MIC | 200 | 6.25 | 50 | 100 | 6.25 |
|  | Pathfinder | **\*400** | 6.25 | **\*100** | 100 | **\*3** |
| IpD03 | MIC | 800 | 6.25 | 25 | 100 | 6.25 |
|  | Pathfinder | **\*6400** | 6.25 | **\*100** | 100 | **\*3** |

<sup>a</sup>All concentrations are provided in µg/mL. Starred (\*) and bolded values indicate when concentrations different from the MIC were used for Pathfinder conjugation assays.

As a basic test for functionality of the engineered constructs in the uncharacterized *Orbaceae* strains, we measured fluorescence levels from pSL1-GFP conjugants. We observed significant variation in GFP fluorescence among these strains (*F*_4,40_ = 322.8, *p* < 0.0001). In particular, lpD01 had such low GFP signal that it was difficult to detect by eye that it fluoresced on a blue light transilluminator (Fig. 2D), but even this weak GFP signal was sufficient for further studies of this strain (see below).

### *D. melanogaster* colonization by engineered *Orbaceae*

We next used our engineered strains to determine if fly-associated *Orbaceae* can colonize *D. melanogaster*. We colonized conventionally reared Canton-S flies using a method we refer to as arena inoculation, in which flies were kept in a container along with an agar plate of fly diet grown with a lawn of one of our strains (see Methods for additional details). We expected that flies would ingest the bacteria while feeding. After 24 hours of inoculation, we transferred flies to fresh diet every 24 hours to eliminate bacteria that survived on the diet rather than within the flies. To ascertain whether bacteria could persist by replicating on the diet itself, we confirmed lack of growth on the yeast-glucose agar fly diet. No growth was observed on the fly diet for strains lpD01, lpD02, and lpD03. BiB and *O. hercynius* CN3 had light growth after 4-5 days, which could potentially be a source of fly recolonization after the initial inoculation arena. Because we did not perform our inoculation with germ-free flies, we anticipated that other microbes in the fly gut might complicate our colonization experiment. However, a benefit to performing our assay with engineered fluorescent bacteria that carry an antibiotic resistance marker is that we can easily identify and select for our strain of interest within a microbiome containing other microbes.

At several time points after inoculation, we washed and crushed 5-6 flies and plated them on selective media with kanamycin. We found that each of these strains can colonize flies to some extent, in contrast to the bee-associated *Orbaceae, Gilliamella apicola* M1-2G, which was not retained at any time point (Fig. 3). Colonization of the fly-derived strains varied between time points and between individual flies in these initial tests. The most consistent findings were that lpD01 was able to robustly colonize flies at every time point, while lpD03 and BiB were lost after day 2 and 4, respectively.

**FIG 3.**
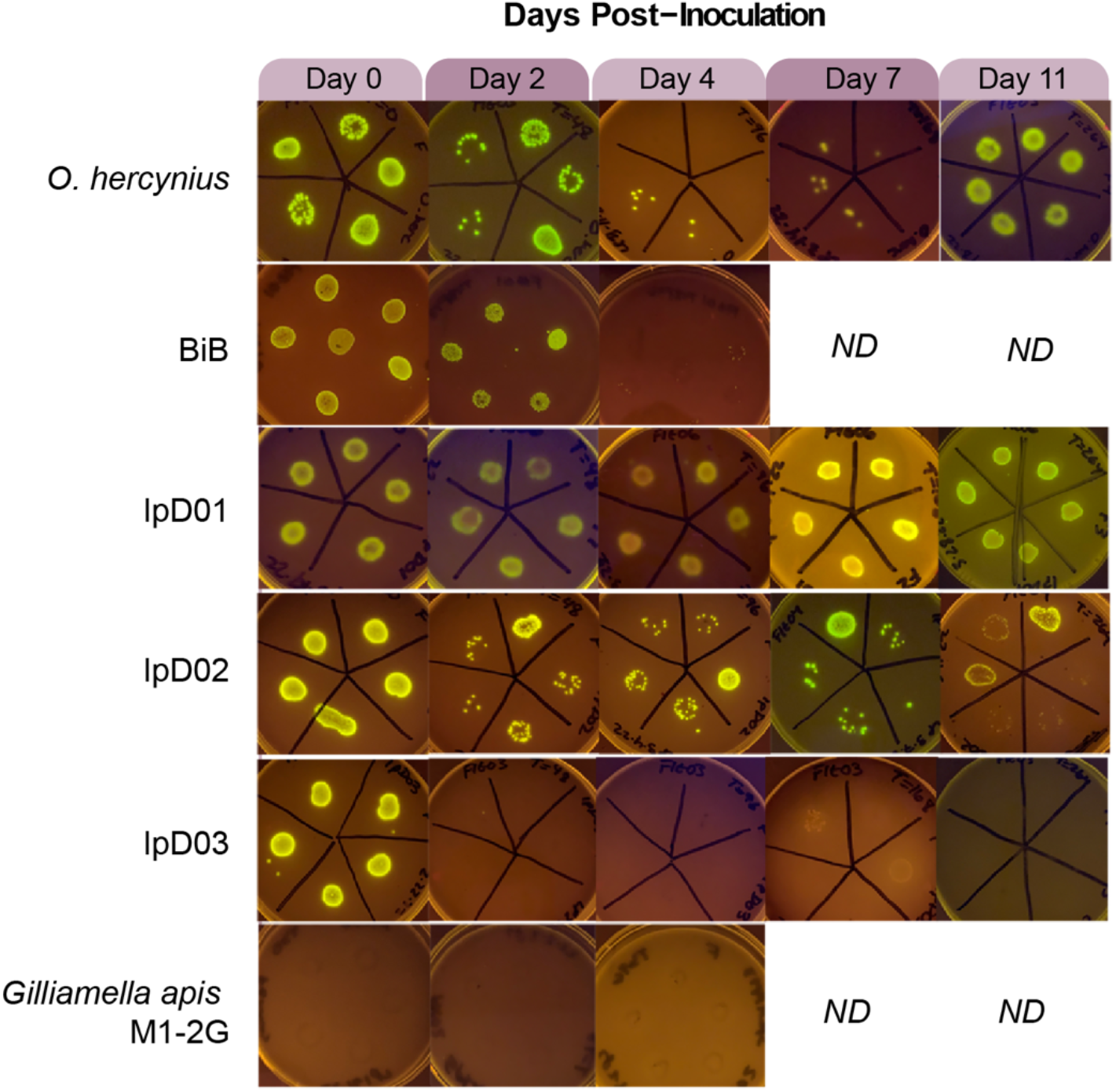
Engineered *Orbaceae* strains can colonize *D. melanogaster*. Images were taken of bacterial growth on selective media containing kanamycin following the growth of bacteria from crushed flies in the days following inoculation. Ability to colonize the host is determined by the presence of many GFP-expressing colonies. Days on which we were unable to collect data or stopped the experiment are indicated by “ND”.

Based on these preliminary results, we decided to track bacterial titer over time in flies colonized with lpD02 and lpD01. To account for variation among experimental populations of flies, we inoculated three separate arenas of flies per trial. The overall trend for lpD02 colonization follows the pattern observed in the qualitative experiment. Over time, the number of colony-forming units (CFU) per fly gradually decreases until day 11, when most flies are no longer colonized (Fig. 4A). However, for lpD01, the average CFU in each arena decreases then increases almost to the initial level seen on day 0 (Fig. 4B). Between day 0 and day 4, average CFU drops (arena 1, *p* = 0.001, arena 2, *p* = 0.0045, arena 3, *p* < 0.0001, Dunn’s test with Bonferroni correction), and most sampled flies were uncolonized on day 4 in arenas 1 and 3. The CFU per fly then increases between day 4 and day 11 for arenas 1 and 3 (*p* = 0.0022 and *p* = 0.0001, respectively, Dunn’s test with Bonferroni correction). The increase between day 4 and day 11 was not significant for arena 2 (*p* = 1, Dunn’s test with Bonferroni correction), but appears to show a similar trend with a slight temporal delay.

**FIG 4.**
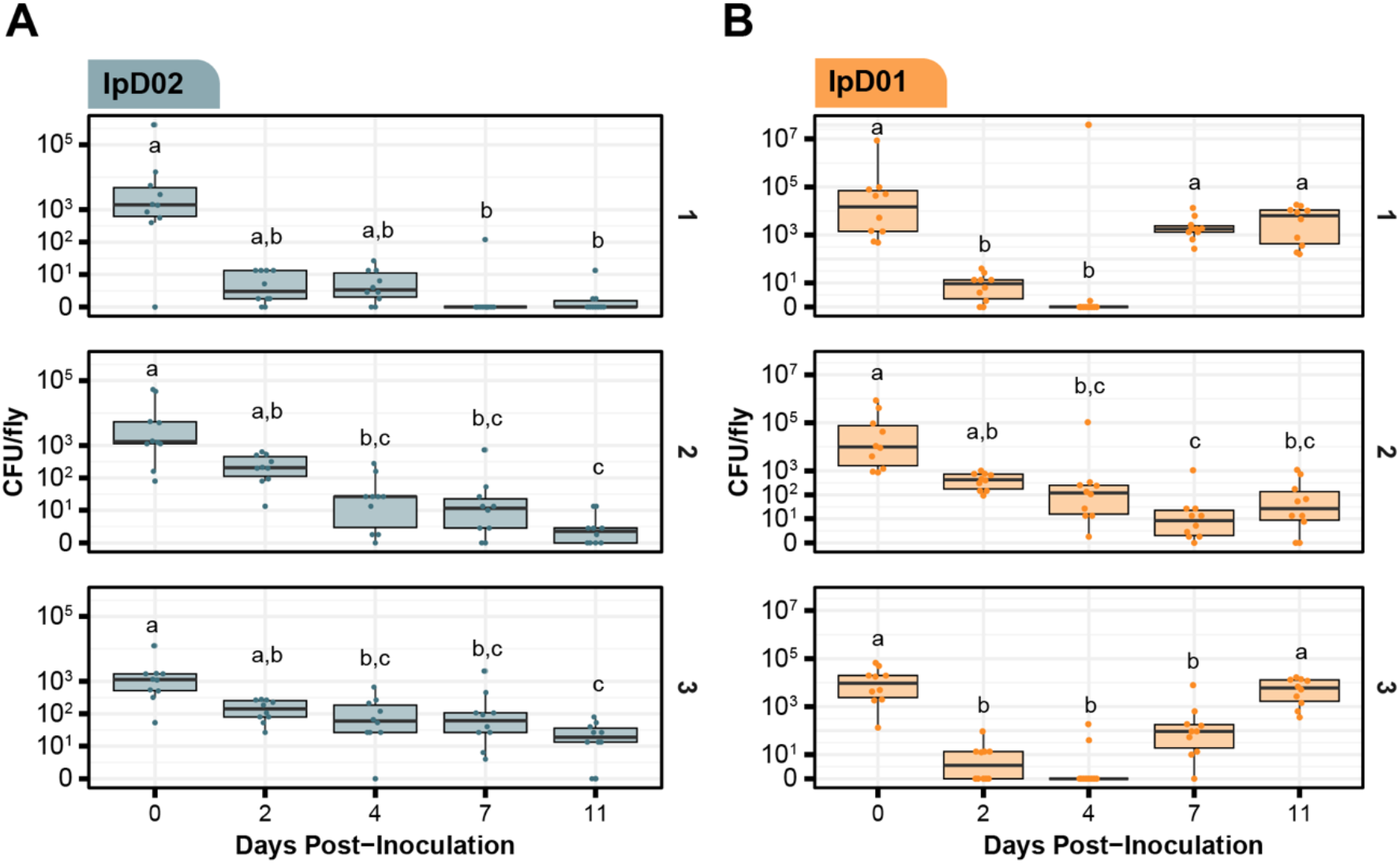
Dynamics of *D. melanogaster* colonization by two newly isolated *Orbaceae* strains. CFU/fly was measured following inoculation with either (A) lpD02 or (B) lpD01. CFUs in each of ten flies per arena were measured at each time point. Data are plotted on a pseudo-log scale so the full range of colonization levels can be shown. Results from three independent arenas are shown in subpanels labeled 1, 2, and 3. Letters above each boxplot represent groupings that are significantly different from one another (*p* < 0.05, Dunn’s test with Bonferroni correction).

### Visualizing lpD01 in the gut of *D. melanogaster*

Based on the results of the bacterial titer assay, we selected lpD01 to visualize colonization of the fly gut by *Orbaceae*. We inoculated 100 flies with the engineered lpD01 + pSL1-GFP strain and reared them for 11 days on fresh diet, replicating the quantification experiment. After this point we dissected the *D. melanogaster* gut and used confocal fluorescence microscopy to assess bacterial localization (Fig. 5). We observed the presence of fluorescent lpD01 in the proventriculus (cardia) of every imaged fly (Fig. 5C, D), and for 2 out of 5 flies lpD01 could also be found colonizing the crop (Fig. 5G, H). These locations are consistent with where gut-associated *Acetobacteraceae* and *Lactobacillus* strains have been observed in *D. melanogaster* (40–42). Bacterial aggregates were present in both crop and cardia (Fig. 5E), suggesting active replication. Throughout the remainder of the gut, we occasionally observed fluorescent cells (Fig. 5J, K), but lpD01 did not robustly colonize the midgut or hindgut regions.

**FIG 5.**
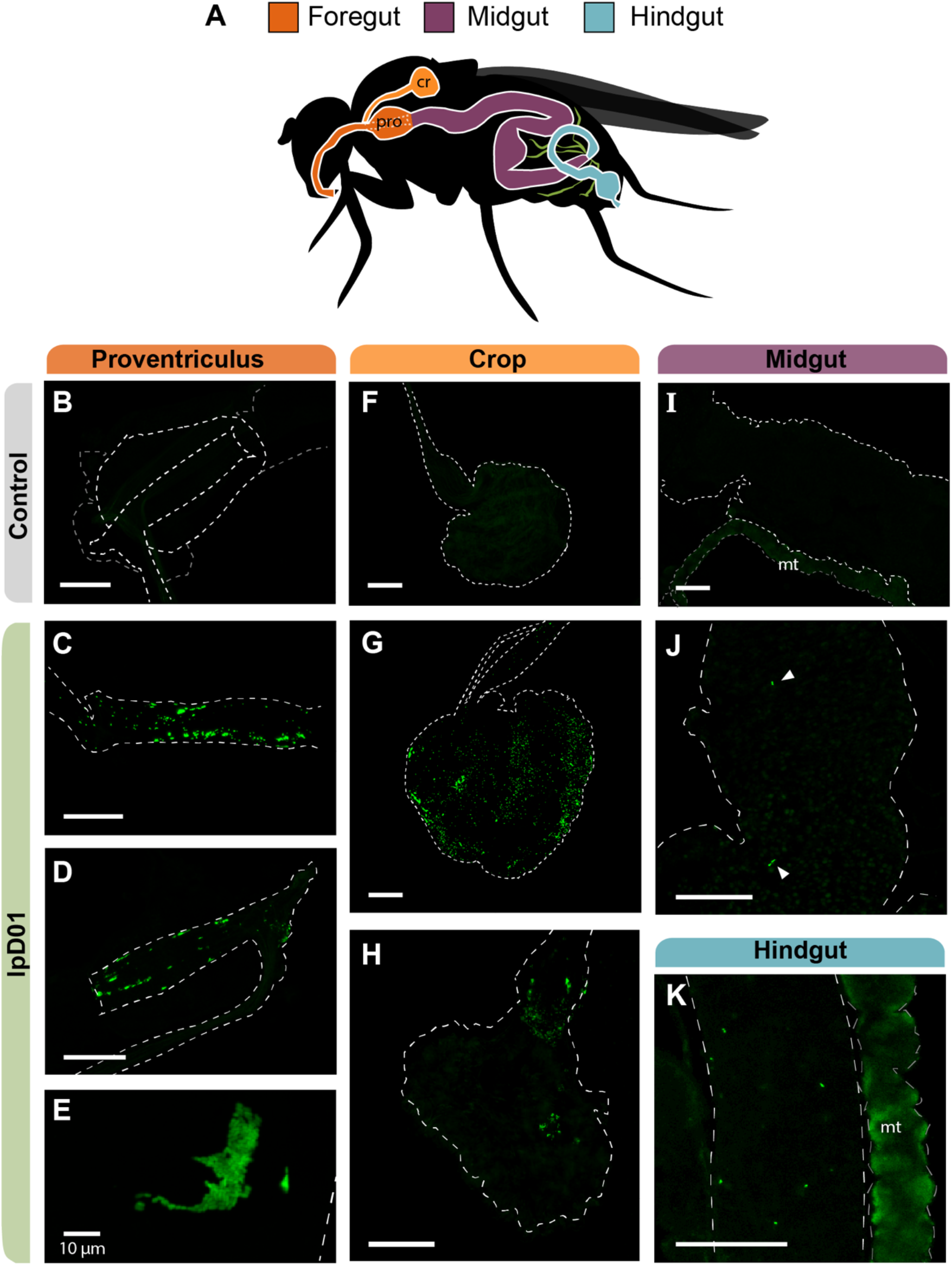
Confocal microscopy of the *D. melanogaster* gut colonized with fluorescent lpD01. (A) Schematic of the *Drosophila* gut with colors designating three regions, foregut, midgut, and hindgut. The crop (cr) and proventriculus (pro) are labelled due to their relevance for the localization of the strain. Malpighian tubules (mt) are depicted in green. (B-K) Confocal images of the dissected gut of one uncolonized (B, F, I) and three colonized flies. Images (C, G, J and K) are from the same fly, (D, H) are from a second fly, and (E) is a from a third. Images (B-D, F-K) were captured at 10× magnification, and image (E) was captured at 40×. Outlines for each of the relevant gut regions are represented by white dashed lines and arrows were added to (J) to point out individual bacterial cells. The Malpighian tubules (mt) exhibited autofluorescence in all flies. GFP intensities were linearly adjusted in each image to highlight bacterial localization. The scale bar for each image represents 100 μm except where indicated.

## DISCUSSION

The Pathfinder plasmid system provides a simple toolkit with a straightforward methodology for genetically modifying non-model bacteria. With this kit we successfully engineered several *Orbaceae* strains isolated from wild flies and utilized their GFP expression and antibiotic resistance cassettes to facilitate colonization experiments in *D. melanogaster*. To our knowledge, our study is the first to demonstrate that *D. melanogaster* can be experimentally colonized with natural fly symbionts from the *Orbaceae* family.

Our results illustrate a common obstacle in synthetic biology: not all plasmid components work well in all bacteria (12). For example, we did not recover *Orbaceae* transconjugants carrying either the RP4 or pBBR1 origins, despite reports that these origins function in a wide range of bacteria in other studies (13, 43). In terms of antibiotic resistance cassette compatibility, our results also matched our expectation that each resistance cassette would not function in every strain. The incompatible strains were mostly those with intrinsic resistance to specific antibiotics. However, the basis of the incompatibility of the Tet resistance cassette with all *Orbaceae* tested is unclear since most were sensitive to low levels of Tet. Possibly, the Tet resistance gene on the Pathfinder plasmids does not express or function effectively in *Orbaceae*. Our varied results highlight the utility of widely surveying for plasmid component functionality when first working with a new strain. The information from Pathfinder informs the selection of plasmids for future experiments.

Many other genetic toolkits have been developed with a similar goal of engineering wild bacteria (9, 16, 44), and Pathfinder has fewer components than other kits. We prioritized building out the complete antibiotic set with RSF1010 because of this origin of replication’s wide compatibility with different bacteria (45, 46). RSF1010 worked well in the *Orbaceae* strains, but it may not replicate in other bacteria. The host ranges of many plasmid origins of replication that function in non-model bacterial species have not yet been exhaustively surveyed. In the future, the kit might be improved by including additional plasmid origins and pairing each of these with the entire antibiotic resistance set. The current Pathfinder toolkit also relies on a single promoter to drive all reporter genes. One could expand the combinations to include different promoters to rapidly survey their functions in a new bacterium, for example by performing multiplex conjugation and then picking the most highly fluorescent or colored colonies. Combinatorial Golden Gate assembly schemes could be used to create sets of plasmids with new combinations of these components.

The Pathfinder kit is limited to plasmid-based expression systems. Plasmid transformation is a common first step in engineering a new bacterium, but it is not ideal for ensuring the long-term stability of engineered constructs. Multicopy plasmids can be especially burdensome when their gene products divert resources from host cell replication (47, 48). Because of this, plasmid-based systems tend to be more likely to rapidly lose function due to the takeover of cells with mutated plasmids that alleviate this burden by inactivating engineered functions, as compared to systems engineered into the chromosome (49). Another potential complication is rapid plasmid loss from a cell population due to segregation in the absence of antibiotic selection. Our colonization experiments in *D. melanogaster* used constant antibiotic selection to prevent plasmid loss. However, administering sufficient levels of antibiotics for selection may be challenging in other environments. Tools designed for chromosomal integration such as transposon systems may be a better option for researchers with these concerns (44). Conjugation has been used to deliver DNA to engineer bacteria *in situ* in gut and soil communities (2, 16, 50). The Pathfinder plasmids are compatible with this approach and could potentially be used to engineer bacteria that are currently unculturable outside of their hosts (1, 51).

The fluorescent *Orbaceae* strains that we built enabled us to easily screen for effective colonization of *D. melanogaster*. We observed differences among isolates in their abilities to colonize flies, as well as variable colonization levels among individual flies inoculated with the same isolate and between the preliminary trials and follow-up experiments with the lpD01 and lpD02 isolates. We colonized non-axenic flies to emulate invasion conditions in which other microbes are already present in the fly gut (40). Differences in the established gut communities of the cohorts of flies that we used could explain some of the variation in our results (41). Using germ-free flies would eliminate these effects (52). Since our flies were housed in a laboratory setting, their diets and microbiomes do not necessarily reflect wild conditions (40), which could also impact the success of *Orbaceae* relative to other gut-associated species of bacteria.

We observed fluorescent *Orbaceae* cells in the crop and proventriculus regions of the foregut in a majority of the flies we colonized with lpD01. The *Drosophila* foregut tends to be more hospitable for bacterial colonization than the midgut, which has a lower pH and undergoes peristalsis along with continual turnover of the peritrophic membrane (21, 41). Accordingly, we observed very few lpD01 cells in the midgut, and other stable colonizers of the *D. melanogaster* gut like *Lactobacillus plantarum* and *Acetobacter thailandicus* also principally localize to the crop and proventriculus (40, 41). Our current results do not reveal whether lpD01 is attaching to and forming biofilms in the gut as has been observed for *L. plantarum* (41).

*Orbaceae* are widespread insect symbionts, but their roles in host biology and the reasons for their host specificity are largely unexplored. *D. melanogaster* offers sophisticated genetic resources for understanding the host side of these microbiome interactions. Our results show that it is also possible to genetically modify *Orbaceae* to begin to dissect these relationships. Because the Pathfinder plasmids are compatible with established Golden Gate assembly schemes and parts libraries (6, 35), they can be used to build new constructs, including systems for knocking out or inserting genes into the bacterial chromosome (6, 44, 53). Such tools would facilitate future studies of insect-*Orbaceae* interactions. It may also be possible to use these genetic tools to control insects that are agricultural pests (1, 54), by isolating and engineering *Orbaceae* native to the tephritid fruit fly *Bactrocera dorsalis* (33), for example.

## MATERIALS AND METHODS

### Isolation of bmE01 from leafhoppers

*Empoasca* sp. leafhoppers were collected by sweep netting *Salvia* sp. plants on the University of Texas campus in Austin, TX (30.289160, –97.738927). Individual leafhoppers were washed by soaking in 70% ethanol for one minute, followed by another minute in 10% bleach. Next, each leafhopper was rinsed 3 times with sterile water and then crushed in 100 μL of sterile saline. Then, 50 μL of 5 different leafhopper samples were plated onto separate Brain Heart Infusion (BHI) agar containing cycloheximide at 100 μg/mL. Colonies were picked and identified based on PCR and Sanger sequencing of the 16S rRNA gene with primers 16SA1F (5’-AGAGTTTGATCMTGGCTCAG) and 16SB1R (5’-TACGGYTACCTTGTTACGACTT) (55). The bme01 isolate studied here was predicted with high confidence to be a *Rosenbergiella* species by both the Ribosomal Database Project classifier tool (56), and BLAST searches of the NCBI 16S ribosomal RNA sequence database (57).

### Isolation of BiB, lpD01, lpD02, and lpD03 from flies

Wild flies were collected at the Brackenridge Field Laboratory in Austin, TX (30.284326, –97.778522). Traps were prepared by adding fermented banana, yeast, and twigs to punctured plastic bottles. Traps were hung from trees for 7 days, and flies were collected each day. Flies were placed on ice or at 4°C immediately after collection and processed within 24 hours.

Flies were immobilized on ice and photographed for morphological identification. They were then washed with 10% bleach to remove surface microbes for 1 minute followed by rinsing in sterile water for 1 minute to remove residual bleach. Legs and wings were removed and placed in 95% ethanol to preserve host DNA. Next, each fly was placed in 50 μl of Insectagro DS2 insect growth medium (IGM) (Corning, VA, USA) and homogenized using a sterile plastic pestle. Dilutions of homogenate were plated on heart infusion agar (HIA) with 5% sheep’s blood. Agar plates were initially incubated at 37°C and 5% CO_2_ (*e.g*., in the case of BiB), but 30°C was later used due to superior growth (*e.g*., in the case of lpD01). Clear or off-white and slower-growing colonies were passaged onto fresh plates multiple times to obtain pure cultures.

To identify the *Orbaceae* bacteria, we amplified and Sanger sequenced the 16S rRNA gene using 27F (5ʹ-AGAGTTTGATCMTGGCTCAG) and *Orbaceae*-specific primer Orb742R (5ʹ-ATCTCAGCGTCAGTATCTGTCCAGAA). Host insects were identified both by morphology and by sequencing PCR amplicons of the barcode region of the COI gene with primers LCO1490F (5ʹ-GGTCAACAAATCATAAAGATATTGG) and HCO2198R (5ʹ-TAAACTTCAGGGTGACCAAAAAATCA) (58). The phylogenetic tree was assembled from 16S ribosomal RNA gene sequences. 16S rRNA gene sequences were aligned in Geneious using MUSCLE (59), and all sites containing ≥50% gaps were stripped. This masked alignment used to infer a phylogenetic tree using IQ-TREE with default options and nonparametric bootstrapping (60). The tree was visualized using iTOL (v5) (61).

### Growth and maintenance of bacterial strains

*Escherichia coli* DH5α, *E. coli* MFD*pir*, and *Sodalis praecaptivus* HS^T^ were grown in LB broth and on LB agar at 37°C. Media was supplemented with 0.3 mM diaminopimelic acid (DAP) for MFD*pir* growth. Following isolation, *Rosenbergiella* was grown at 30°C on BHI broth or agar. *Orbus hercynius* CN3 was acquired from the German Collection of Microorganisms and Cell Cultures (DSMZ) (DSM 22228). Following isolation, *Orbaceae* strains were determined to be culturable in either IGM or BHI broth, and on BHI agar or HIA with or without 5% defibrinated sheep’s blood. For robust growth, BHI and HIA + 5% sheep’s blood were preferred, but media were selected based on the needs of the assay. All fly *Orbaceae* were grown at 30°C with 5% CO_2_. *Gilliamella apis* M1-2G was grown on HIA + 5% sheep’s blood at 35°C with 5% CO_2_. Antibiotic concentrations used in this study are shown in Table 2.

### MIC tests to determine antibiotic susceptibility

We performed MIC assays to determine the appropriate selective antibiotic concentration for each bacterial strain. To do so, we prepared 96-well plates with 2-fold dilutions of each antibiotic, ranging from 400 – 6.25 μg/mL in 100 μL media. One microliter of each strain was inoculated in triplicate for each condition. After allowing the strains to grow for 1-3 days, the plates were inspected visually to determine the minimum inhibitory concentration for each antibiotic. These MIC values were used to guide how much antibiotic was used for selection during the Pathfinder conjugation process (Table 2).

### Assembly of the Pathfinder plasmids

All cloning procedures were carried out in *E. coli* strain NEB5alpha (#C2987H, New England Biolabs) cultured overnight aerobically at 37**°**C in LB broth or solid LB agar. Antibiotics were supplemented when necessary for plasmid selection or maintenance at the following concentrations: Kanamycin (Kan) (50 μg/mL), Spectinomycin (Spec) (60 μg/mL), Gentamicin (Gent) (25 μg/mL), Chloramphenicol (Cam) (20 μg/mL), Erythromycin (Ery) (250 μg/mL), Tetracycline (Tet) (10 μg/mL), and Carbenicillin (Carb) (100 μg/mL).

We designed the Pathfinder plasmid series to have a variety of broad-host-range origins, antibiotic resistance genes, and highly expressed visible reporters suitable for rapid identification and testing in newly isolated bacteria. We started with pBTK402, a broad-host-range plasmid with a RSF1010 origin that we previously engineered to remove any BsaI and BsmBI cut sites and make it suitable for Golden Gate Assembly (6). The pBTK402 plasmid was designed to function as a Type 8 dropout vector for the BTK Golden Gate assembly scheme, and all of the main Pathfinder plasmids retain this attribute. We replaced the weakly expressed *rfp* on pBTK402 with a visible red chromoprotein (RCP) (Bba_E1010) expressed from the strong CP25 promoter and associated ribosome binding site (RBS) from plasmid pBTK569. This promoter-RBS combination appears to lead to robust protein expression in Proteobacteria. This plasmid was re-designated “pSL1” and all subsequent pSL plasmids are derived from pSL1. Plasmid pSL1-GFP replaces the RCP reporter with GFP. Plasmids pSL2 – pSL7 were constructed by replacing the Kanamycin resistance allele (*aphA-1*, Kan^R^) present in pSL1 with an alternate antibiotic resistance allele and associated upstream promoter. pSL9 and pSL25 replace the RSF1010/RCP origin and reporter with pBBR1/E2-Crimson (E2C) and RP4/blue chromoprotein (BCP), respectively (see Table 1).

To construct plasmids, we first designed assembly primers using Benchling (http://www.benchling.com) and added Golden Gate-compatible BsmBI cut sites. We ordered DNA primers from Integrated DNA Technologies and then amplified PCR products using either Q5 Hot-Start Master Mix (#M0494S, New England Biolabs) or KOD XL (#71087-3, Millipore Sigma) according to manufacturer’s instructions. We purified PCR products with a QIAquick PCR Purification Kit (#28104, QIAGEN), assembled them using a NEBridge Golden Gate Assembly Kit (BsmBI-v2) (#E1602S, New England Biolabs), and then electroporated 1 μL of the reaction into electrocompetent NEB5alpha. Cells recovered for 1 hour and were then plated on appropriate selective media. Plasmids were initially verified by Sanger sequencing of the assembly junctions and later by whole-plasmid Illumina sequencing on an iSeq 100. Three plasmids (pSL2, pSL4, and pSL5) contain a duplication of a BsaI restriction site and the adjacent CP25 promoter and ribosome binding site for RCP. Whether this duplication affects RCP expression is unknown. These plasmids still function as Type 8 dropout vectors for Golden Gate assembly.

We next transformed all pSL plasmids into the DAP auxotrophic conjugation donor strain *E. coli* MFD*pir* (36). We could not successfully transform pSL5, however, due to intrinsic erythromycin resistance in MFD*pir*. These strains of MFD*pir* were used individually or combined for subsequent multiplex conjugation assays. A preliminary study examined various aspects of how the RSF1010-based Pathfinder plasmids function in *E. coli*, including how different antibiotic markers and concentrations affect plasmid copy number and reporter output and how stably they are maintained over many serial transfers in laboratory cultures with or without antibiotic selection (62).

### Conjugation of the Pathfinder plasmids into insect associated bacteria

We created the Pathfinder conjugation mix by first growing up each of the donor strains separately to saturation. At this point we measured the optical density at 600 nm (OD600) of each strain and resuspended it at an OD600 of 1. Equal volumes of each strain were combined into a single tube along with 16% glycerol then distributed into PCR tubes and frozen at −80°C. For conjugations, this mix can be thawed and added straight to the first conjugation step.

To perform the conjugation itself, we started with 1 mL of an overnight culture of the target bacteria. The culture was pelleted (1000 × *g* for 6 minutes) and washed once with 145 mM NaCl saline to remove any residual media, and then resuspended in saline to OD600 = 1. Twenty-five μL of the sample was combined with 25 μL of the thawed Pathfinder plasmid mix and spot-plated on media compatible for the growth of both *E. coli* and *Orbaceae* spp. (BHI for all strains except lpD02, which was grown on BHI + 5% sheep blood) plus DAP. After 1-2 days of growth, we scraped up all the growth from the conjugation spot and suspended it in 1 mL of saline. This was washed and resuspended twice with 1 mL sterile saline to remove residual DAP. The resuspended sample was divided into 5 equal portions and serially diluted 10-fold to a final dilution of 1 × 10^−5^. To plate these dilutions, 5 μL of each replicate and dilution were spotted onto antibiotic plates at each bacterium’s MIC, along with a zero-antibiotic control. Plates were left to dry and then placed in an incubator at the optimal temperature for each bacterium. If one of the antibiotic concentrations proved to be too low or too high following conjugation, it was adjusted accordingly, and the procedure was repeated (Table 2). To confirm successful conjugation, colonies were first visually examined for expression of fluorescent reporter genes. Then, one colony from each antibiotic condition was regrown in a liquid medium, and a dilution of this culture was used as template for whole-cell PCR and 16S rRNA gene sequencing with primers 16SA1F and 16SB1R, as described above. Colonies were counted and the efficiency of conjugation was determined relative to growth on the zero-antibiotic control plate.

### Imaging bacterial colonies

To ensure accurate categorization of each of the fluorescent reporters in strains where the fluorescence was not as bright, we visualized plates on a Typhoon 9500 imager (GE Healthcare Bio-Sciences, Uppsala, Sweden). To visualize RCP fluorescence, we imaged with 532 nm excitation and 575 nm emission. To distinguish E2C fluorescence from RCP, we imaged the plate with 635 nm excitation and 665 nm emission. Images were processed and counted in Fiji (version 1.53q) (63).

### Measuring GFP expression in different bacteria

To measure the level of GFP expression, we assessed three separate colonies picked from the Pathfinder conjugation plate. Each colony was first grown to saturation, then re-grown with a starting OD600 of 0.05 in 5 mL Insectagro DS2 in a test tube. After re-growth, 1 mL of each culture was pelleted and resuspended in 600 μL saline. OD600 and GFP expression (485 nm excitation, 535 nm emission) were measured for 200 μL of each resuspension in triplicate using a Tecan Infinite 200 Pro plate reader. GFP expression was normalized to the OD600 measurement for each sample.

### Growth and maintenance of *Drosophila* stocks

*D. melanogaster* Canton-S were acquired from the Bloomington Drosophila Stock Center (Bloomington, IN). Fly stocks were reared on Formula 4-24 Instant Drosophila Medium (Carolina Biological Supply Company, Burlington, NC). For experiments, stocks were swapped to a yeast-glucose agar (YGA) diet containing brewer’s yeast, D-glucose, agar, and water (52). Nystatin (10 μg/mL) and kanamycin (10 μg/mL) were added to the diet where specified to prevent the growth of fungal contaminants and ensure the maintenance of engineered strains, respectively. Stocks were maintained at 25°C during experiments, with a 12L:12D photoperiod in Percival I-36LLVL incubators (Perry, IA, USA).

### Inoculation of *Drosophila* with *Orbaceae* strains

*D. melanogaster* was inoculated with the engineered *Orbaceae* using a method we refer to as arena inoculation. For these experiments, adult female *D. melanogaster* were first removed from the commercial diet and transferred to YGA diet for 48 hours to clear their gut from the preservatives in the commercial diet. In the meantime, HIA + 5% sheep blood + Kan plates were inoculated with 100 μL of OD600 = 1 *Orbaceae* + pSL1-GFP and allowed to grow for 1-2 days as needed. The bacterial plate was then taped to the bottom of a lidded 12oz SelecTE plastic container “arena” (Berry Plastics, Evansville, IN) modified with a mesh vent on its lid and a small port for administering CO2. YGA flies were added to an arena and allowed to feed for 24 hours. After this inoculation step, flies were transferred daily to fresh YGA + nystatin + Kan diet tubes.

### Quantification of bacterial colonization

Each bacterial strain was first delivered to female adult flies 3-5 days post-eclosion using the methodology described above. For this set of experiments, 3 different arenas were used to inoculate 3 separate populations of female *D. melanogaster*. Equal numbers of flies were placed into each arena, ranging from 100-150 in each arena for each round of this experiment. After inoculation, each separate population of flies was transferred to their own tube of YGA + nystatin + Kan diet. Flies were transferred to fresh diet daily. At day(s) 0, 2, 4, 7, and 11 post-inoculation, 10 flies from each independent population (30 total) were crushed and plated to determine their quantity of bacterial colonization.

To prepare samples for crushing, we first washed them in 10% bleach to ensure that any bacteria plated would not be from the outside of the fly body. Flies were immobilized for this procedure by first placing them at −20°C for 1 minute, and then keeping the tubes on ice during the washes. They were soaked in 500 μL 10% bleach for 1 minute, washed with 500 μL 1× phosphate buffered saline (PBS), then resuspended in 200ul PBS and homogenized with a sterile plastic micropestle. A set of 1:10 serial dilutions in PBS were then carried out for each sample. Five microliters of each dilution were spotted onto agar plates (HIA + 5% sheep blood, Kan 50 μg/mL, nystatin 10 μg/mL) in triplicate. The remaining homogenized fly mixture was then centrifuged at 1000 ×*g* for 6 minutes to pellet the remaining bacteria. The pellet was resuspended in 15 μL saline and plated to ensure the detection of bacteria that were present in low abundance.

### Imaging *Drosophila* guts with confocal microscopy

Fifty female *D. melanogaster* were colonized with lpD01 + pSL1-GFP in two separate arenas and maintained for 11 days along with an uncolonized control population. To ensure that flies were colonized with the bacteria, 3 flies from each population were crushed and plated on day 9 as described in the prior methods section. On day 11, all flies were transferred to a tube without diet for 12-18 hours to empty the gut of any diet-associated particles that may complicate imaging. The entire gut of a selection of the flies was dissected and mounted in PBS on a glass slide. The dissected guts were imaged using a Leica MZ16 Fluorescent Stereoscope in the GFP channel to visualize bacterial colonization. Images were linearly adjusted to highlight the bacterial localization using Fiji software (version 1.53q) (63).

## Data availability

The data that support the findings of this study and newly isolated bacteria reported here are available from the authors upon reasonable request. The Pathfinder plasmids have been deposited in Addgene (Table 1).

## ACKNOWLEDGEMENTS

We thank Bibiana Toro and other members of the 2018 UT Austin iGEM team for assistance with Pathfinder toolkit development and testing. We thank John McCutcheon for the gift of *Sodalis praecaptivus*. We thank Thilini Wijesekera for providing stocks of *D. melanogaster* Canton-S. This research made use of equipment at the Microscopy and Imaging Facility of the Center for Biomedical Research Support at The University of Texas at Austin (RRID# SCR_021756). Funding was provided by the U.S. Army Research Office (W911NF-20-1-0195) to J.E.B. and N.A.M. and by the U.S. National Institutes of Health (R35GM131738) to N.A.M. K.M.E. acknowledges a UT Austin University Graduate Continuing Fellowship, T.A. and B.M. acknowledge UT Austin TIDES Summer Research Fellowships, and B.M. acknowledges an Army Educational Outreach Program (AEOP) Undergraduate Research Apprenticeship.

